# SWALO: scaffolding with assembly likelihood optimization

**DOI:** 10.1101/081786

**Authors:** Atif Rahman, Lior Pachter

## Abstract

Scaffolding *i.e.* ordering and orienting contigs is an important step in genome assembly. We present a method for scaffolding based on likelihoods of genome assemblies. Generative models for sequencing are used to obtain maximum likelihood estimates of gaps between contigs and to estimate whether linking contigs into scaffolds would lead to an increase in the likelihood of the assembly. We then link contigs if they can be unambiguously joined or if the corresponding increase in likelihood is substantially greater than that of other possible joins of those contigs. The method is implemented in a tool called Swalo with approximations to make it efficient and applicable to large datasets. Analysis on real and simulated datasets reveals that it consistently makes more or similar number of correct joins as other scaffolders while linking very few contigs incorrectly, thus outperforming other scaffolders and demonstrating that substantial improvement in genome assembly may be achieved through the use of statistical models. Swalo is freely available for download at https://atifrahman.github.io/SWALO/.

## Background

The emergence of second generation sequencing technologies [1–4] has led to development of various assays to probe many aspects of interest in molecular and cell biology due to the low cost and high throughput. However, a prerequisite for running many of these assays, genome assembly is yet to be solved adequately using second generation sequencing reads. Although the emergence of long read technologies such as single molecule real time (SMRT) [5] and nanopore sequencing [6] is transforming genome assembly, the high cost, low sequencing coverage and high error rates of these technologies mean that most of the genomes are being assembled from second generation data or a combination of the two. In addition an extensive amount of second generation data exists already that can be better utilized. In this paper, we demonstrate that considerable improvement in genome assembly using second-generation sequencing can be achieved through the application of statistical models for sequencing. The statistical approach we introduce may also be applied to genome assembly using long reads.

Genome assembly typically consists of two major steps. The first step is to merge overlapping reads into contigs which is commonly done using the *de Bruijn* or overlap graphs. In the second step, known as “scaffolding”, contigs are oriented and ordered using paired-end or mate-pair reads (we use the term read pair to refer to either). Scaffolding, introduced in [7], is a critical part of the genome assembly process and has become of increased importance due to short length of second generation reads. It is hence built into most assemblers [8–12] and a number of standalone scaffolders such as Bambus2 [13, 14], MIP [15], Opera [16, 17], SCARPA [18], SOPRA [19], SSPACE [20], BESST [21] have also been developed to better utilize read-pair information and to resolve ambiguities due to repetitive regions. Most of the scaffolding algorithms rely on heuristics or user input to determine parameters such as minimum number of read-pairs linking contigs to join them ignoring contig lengths, sequencing depth and sequencing errors. In an in-depth study, Hunt *et al.* evaluated scaffolding tools on real and simulated data and observed that although many of the scaffolders perform well on simulated datasets, they show inconsistent performance across real datasets and mapping tools [22]. Their results demonstrate that SGA, SOPRA and ABySS are conservative and make very few scaffolding errors while SOAPdenovo identified more joins at the expense of greater number of errors indicating a scaffolding method achieving a better trade-off of the two may be possible.

Here we present a scaffolding method called ‘scaffolding with assembly likelihood optimization (Swalo)’. Swalo learns parameters automatically from the data and is largely free of user parameters making it more consistent than other scaffolders. It is also able to make use of multi-mapped read pairs through probabilistic disambiguation which most other scaffolding tools ignore. The method is grounded in rigorous probabilistic models yet proper approximations make the implementation efficient and applicable to practical datasets. We analyze the performance of Swalo using datasets used by Hunt *et al.* and find that Swalo makes more or similar number of correct joins as other scaffolders while making very few incorrect joins. We also compare Swalo with scaffolding modules built into various assemblers using GAGE datasets [23] and observe that final results obtained by applying Swalo on contigs generated by assemblers are generally better than applying the in-built scaffolding modules of those assemblers.

## Results

### Overview of SWALO

Our scaffolding method called Swalo is based on a generative model for sequencing [24]. Figure 1 illustrates the main steps of Swalo. In the first step, reads are aligned to contigs, the insert size distribution and error parameters are learned using reads that map uniquely and the likelihood of the set of contigs is computed using a generative model. We then construct the bi-directed *scaffold graph* which contains a vertex for each contig and there is an edge between contigs if joining them would result in an increase in the likelihood. It uses probabilistic models to estimate maximum likelihood gaps between contigs correcting for the issue that we may not observe inserts from the entire distribution of insert sizes due to gaps between contigs and lengths of contigs [25, 26]. It then approximates whether joining contigs would result in an increase in the genome assembly likelihood. We use the EM (expectation maximization) [27,28] algorithm to resolve multi-mapped read pairs. Contigs are then joined if the increase in the likelihood is significantly higher than that of all other conflicting joins as determined by a heuristic. Moreover, we select multiple joins consistent with one another using the dynamic programming algorithm for the weighted interval scheduling problem. Each of these steps is described in more detail in the Methods section and in Supplementary Notes 3.1-3.3. Our scaffolding method i) learns parameters from the data making it largely parameter free (Supplementary Note 3.4), ii) makes use of multi-mapped read pairs that are ignored in most scaffolders, and iii) is able to accurately estimate gaps between contigs facilitating gap-filling.

**Figure 1:**
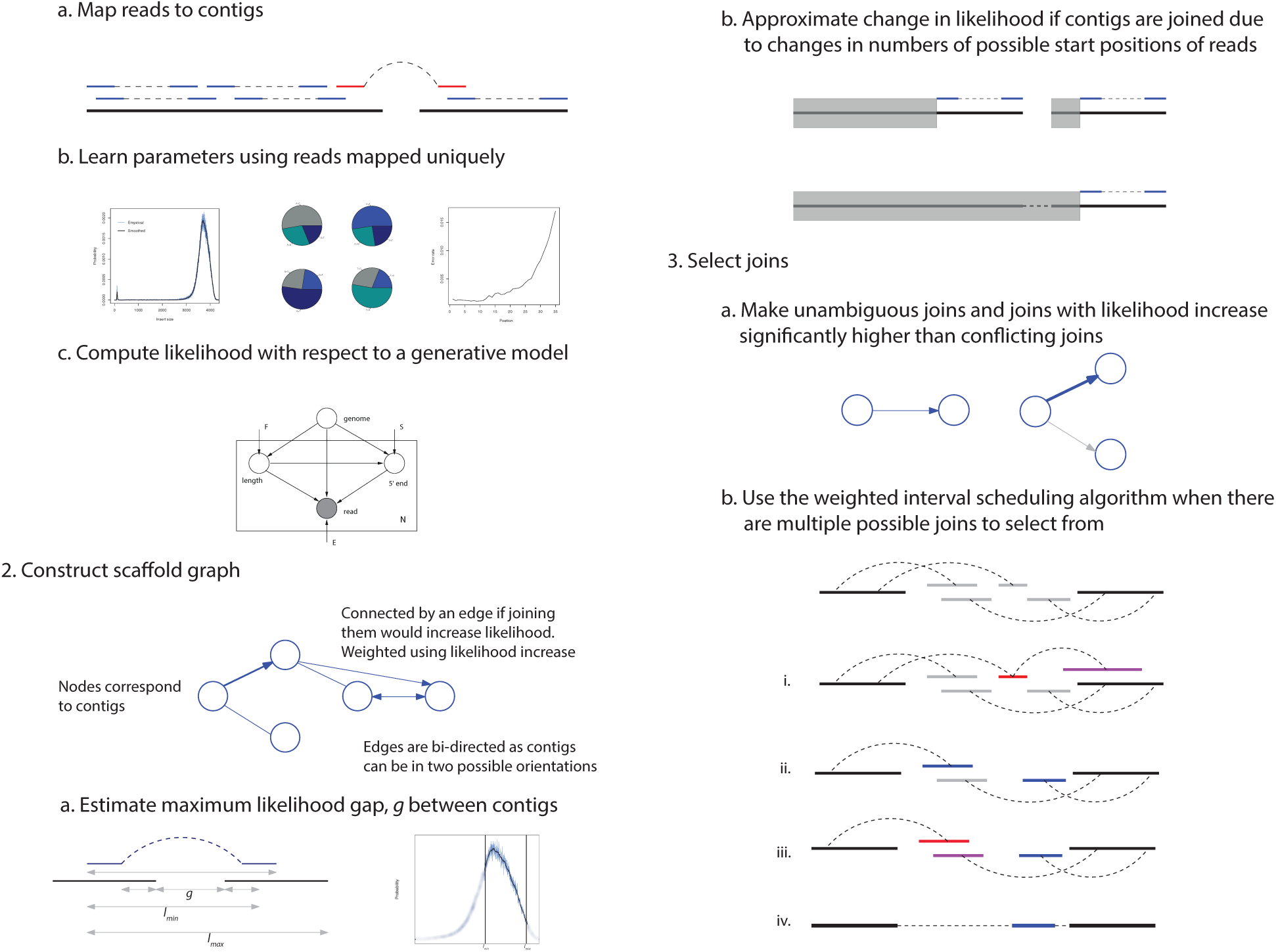
Overview of SWALO. 1. Reads are aligned to contigs, uniquely mapped reads are used to learn the insert size distribution and error parameters, and then the likelihood of of the set of contigs is computed. 2. The scaffold graph is constructed by first estimating maximum likelihood gaps, *g* between contigs using the EM algorithm to resolve multimapped read-pairs taking into account that only inserts of sizes between *l_min_* and *l_max_* will be observed due to gaps between contigs and lengths of contigs, and then approximating whether changes in number of possible start sites of reads (the regions shaded in grey) lead to an increase or decrease in the assembly likelihood. 3. Finally, we make the joins that are unambiguous or correspond to likelihood increase significantly higher than other conflicting joins. If there are multiple contigs (grey) that fit into the gap between contigs being joined we select from them using the dynamic programming algorithm for the weighted interval scheduling problem in following steps, i. Remove contigs (red) with inconsistent edges to other contigs. ii. Select consistent set of contigs (blue) that optimizes likelihood, iii. Remove selected contigs (red) with likelihood increase not significantly higher than conflicting ones not selected (purple), iv. Merge them into scaffolds.

### Comparison with stand-alone scaffolders

To compare performance of Swalo with other stand-alone scaffolders, we use the datasets used by Hunt *et al.* to evaluate scaffolding tools [22]. In addition to the scaffolders considered in the study, we include recently published versions of Opera (Opera-LG [17]) and BESST [29]. The datasets include four simulated datasets from *S. aureus* and six real datasets from *S. aureus, R. sphaeroides, P. falciparum* and human chromosome 14 (Supplementary Table 1). Among these the *S. aureus, R. sphaeroides*, and human chromosome 14 datasets were also part of the GAGE project [23]. The contigs were generated using Velvet [8] which were then aligned to the reference and split at locations of error to ensure misassembly free contigs. Please see [22] for more details on the datasets. We use Bowtie [30] and Bowtie 2 [31] for mapping reads, analyze results using the scripts provided in [22] and when applicable use the same parameter values for mapping and scaffolding as used in the paper by Hunt *et al.* (all parameters used given in Supplementary Table 2). Similar to their paper we use number of correct joins and number of incorrect joins for comparison as contiguity statistics such as N50 scaffold length biases evaluation towards scaffolders making more joins whether they are correct or incorrect while corrected N50 scaffold length leads to favorable assessment of scaffolders with more correct joins even if that is at the expense of many more incorrect joins compared to other scaffolders.

Supplementary Table 3 summarizes performance of scaffolding tools on simulated datasets. We find that Swalo makes no incorrect joins for any of the datasets. For 100kb contigs Swalo was able to make 100% of the correct joins using either library and all aligners. When the insert size library of mean 500bp was used to scaffold 3kb contigs, Swalo made 99.0%, 99.3% and 99.0% correct joins using Bowtie 2, Bowtie with 0 (-v 0) and 3 (-v 3) mismatches respectively. The only scaffolder that makes more than 99.3% correct joins is Opera at 99.8% when used in conjunction with BWA but this is at the cost of making 0.2% incorrect joins. For 3kb contigs and 3kb insert size library, Swalo made 99.6%, 99.8% and 99.6% correct joins for three mapping modes. No other scaffolder made more than 99.6% correct joins. It is worth pointing out that Swalo was able to make more correct joins when used with Bowtie -v 0 compared to Bowtie -v 3 and Bowtie 2 which may be due to reads not being mapped to some regions for the latter two.

Performance of Swalo in comparison to other scaffolders for real datasets is illustrated in Figure 2, Supplementary Figure 2 and Supplementary Tables 4-7. For the *S. aureus* dataset from GAGE, we find that Swalo made more correct joins than all other scaffolders while making 1, 1 and 2 incorrect joins in the three runs corresponding to the three ways of mapping reads. However, closer inspection reveals that one join labeled incorrect in each case is in fact a join from the end to the start of a circular sequence and is actually correct. Similarly for the *R. sphaeroides* dataset, more correct joins are made by Swalo than all other scaffolders when used in conjunction with Bowtie 2. Again 3 joins that are marked as incorrect are joins linking the ends to the starts of circular chromosomes or plasmids. we observe that the sequencing error rate for this dataset is high compared to the *S. aureus* dataset. So, number of reads mapped by Bowtie is quite low [22] resulting in lesser number of joins made by Swalo and other scaffolders when Bowtie is used compared to Bowtie 2.

**Figure 2:**
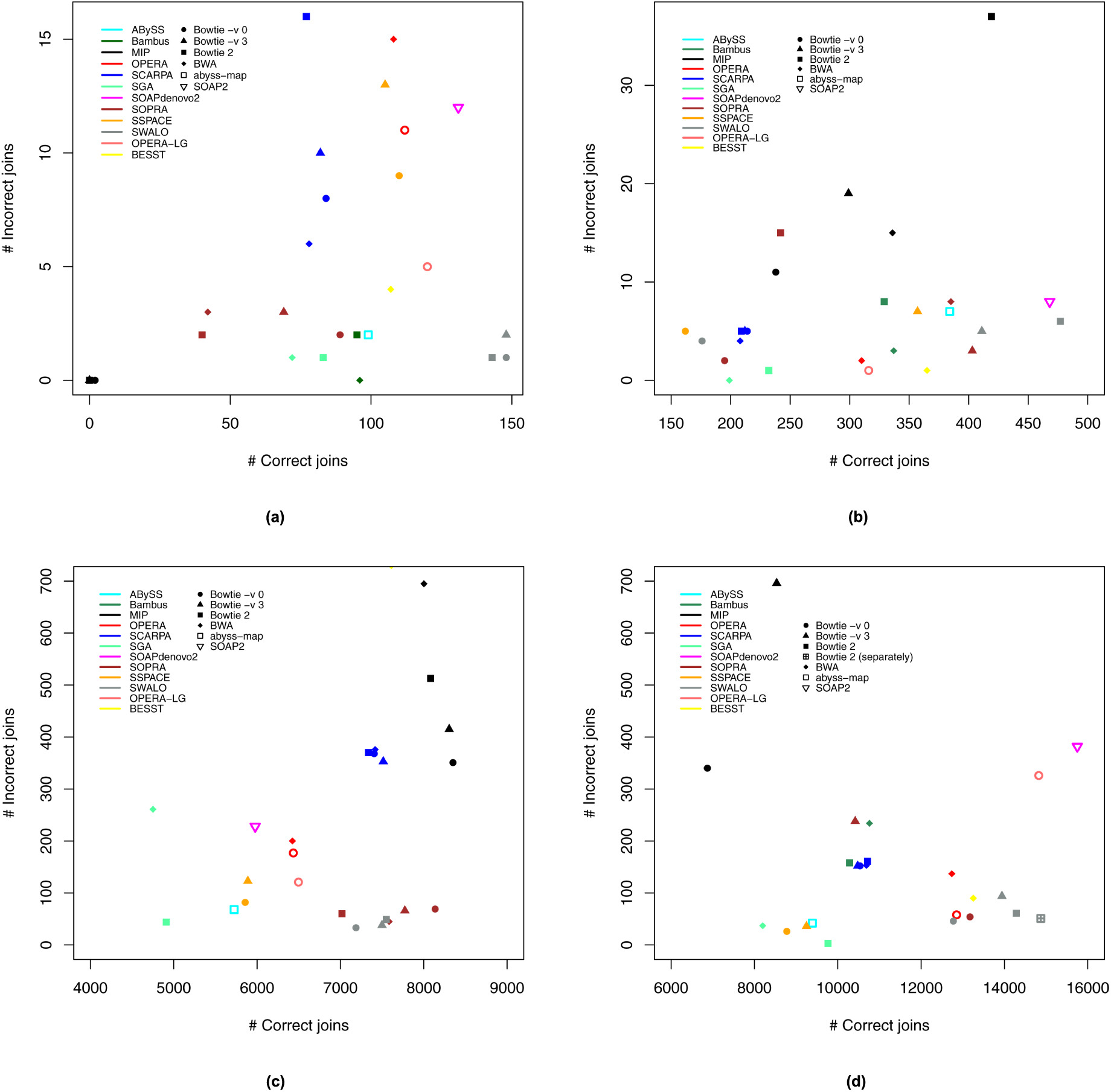
Performance of scaffolders. Scatter plots showing number of correct joins vs incorrect joins made by Swalo and other scaffolders on (a) *S. aureus* data, (b) *R. sphaeroides* data, (c) *P. falciparum* combined short and long insert data, and (d) human chromosome 14 combined long insert and fosmid library data. Up to 1 and 3 joins in (a) and (b) respectively made by Swalo (and possibly other scaffolders) labeled incorrect are joins from end to start of circular sequences and are therefore correct. Values for all scaffolders except Swalo, Opera-LG and BESST are from [22].

The *P. falciparum* genome is known to be hard to assemble due its low GC content. In this case although Swalo does not make more correct joins than all other scaffolders as in other cases, the numbers of correct joins made are only slightly less than that of SOPRA, MIP and SCARPA while number of incorrect joins is less than or similar to what SOPRA made and much less than the numbers for SCARPA and MIP. We observe that many of the contigs have strings of consecutive ‘A’s or ‘T’s where very few reads are mapped to by aligners leading to poor gap estimates which may explain the comparatively smaller numbers of links by Swalo.

Finally, for the combined human chromosome 14 dataset, Swalo makes more correct joins than all other scaffolders except SOAP2 and Opera-LG both of which make more than three times incorrect joins compared to the highest number of incorrect links by Swalo and more than six times the best result by Swalo. Supplementary Figure 1 shows that the long jumping library is in fact a mixture of inserts of two sizes. When they are mapped and used to estimate gaps separately before scaffolding, and the fosmid library is applied on the output, the results improve both in terms of the increase in number of correct joins and decrease in number of incorrect joins.

### Comparison with other scaffolding modules

While Hunt *et al.* performed a comprehensive evaluation of stand-alone scaffolding tools, scaffolding modules built into some assemblers such as ALLPATHS-LG [10,32], MaSuRCA [33], CABOG [34] were left out as they cannot be run independently. In order to assess performance of Swalo in comparison to scaffolding modules of these assemblers, we ran Swalo on the contigs generated by the assemblers obtained from the GAGE project and compared the results with final results of contig assembly and scaffolding by each of these assemblers. The results are shown in Table 1. It reveals that Swalo makes fewest number of incorrect joins in all cases while making more or similar number of correct joins as ALLPATHS-LG and CABOG. For the human chromosome 14 dataset, there are 17 more joins in scaffolds generated by ALLPATHS-LG compared to Swalo output. However, 9 more incorrect joins are made by ALLPATHS-LG than Swalo. Although MaSuRCA makes more correct joins than Swalo, this is at the expense of more incorrect joins which is drastically high for human chromosome 14. It may be noted that information such as actual position of reads in contigs and distance between contigs in assembly graph that were available to the assemblers could only be inferred by Swalo by mapping the reads back to the contigs using a short aligner such as Bowtie. We believe that if these information were made available by assemblers, the results could be further improved.

**Table 1:** Comparison of performance of Swalo with scaffolding modules built into assemblers using GAGE datasets. Comparison of results obtained by running Swalo on contigs generated by various assemblers with final results obtained by those assemblers after scaffolding.

| Dataset | Contig stats | Original scaffold stats |  | SWALO stats |  |
| --- | --- | --- | --- | --- | --- |
| Assembler | Number | Correct | Incorrect | Correct | Incorrect |
| *S. aureus* |  |  |  |  |  |
| ALLPATHS-LG | 60 | 48 | **0** | **49** | **0** |
| MSR-CA | 94 | **74** | 3 | 64 | **2** |
| *R. sphaeroides* |  |  |  |  |  |
| ALLPATHS-LG | 204 | 170 | **0** | **186** | **0** |
| MSR-CA | 395 | **347** | 5 | 228 | **3** |
| CABOG | 322 | 187 | 5 | **239** | **1** |
| Human chromosome 14 |  |  |  |  |  |
| ALLPATHS-LG | 4529 | **4259** | 45 | 4242 | **36** |
| MSR-CA | 30091 | **27521** | 1145 | 20242 | **167** |
| CABOG | 3361 | 2845 | 37 | **2980** | **32** |

### Time and memory requirements

Swalo uses statistical models to estimate gaps between contigs and the change in genome assembly likelihood achieved if contigs are joined. As a result it is more computationally intensive than some other scaffolders. However, we make necessary approximations to make Swalo fast, memory efficient and scalable to large datasets. Figure 3 and Supplementary Table 8 show running times and memory usage of Swalo using 32 cores on a machine with Intel Xeon E5 2.70GHz processors to scaffold Hunt *et al.* datasets.

**Figure 3:**
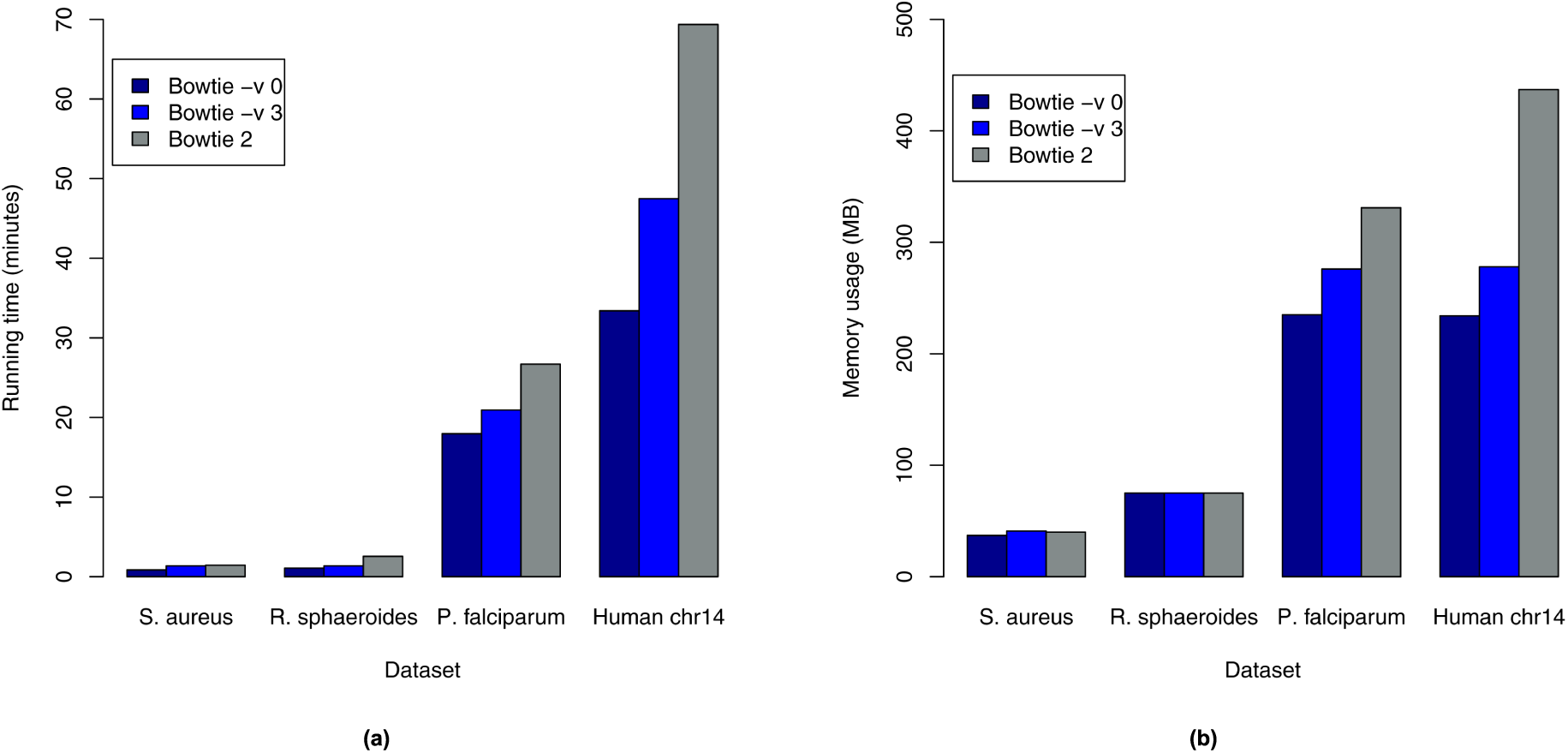
Running time and memory usage of SWALO. Barplots showing (a) running times and (b) memory usage of Swalo using 32 cores on a machine with Intel Xeon E5 2.70GHz processors to scaffold Hunt *et al.* datasets.

Although a comparison of running times is not appropriate since Swalo was run on a different machine to other scaffolding tools, we would like to note that Swalo took from approximately a minute for *S. aureus* datasets to approximately 70 minutes for combined human chromosome 14 dataset to run (excluding the time required for mapping). The memory usage ranges from around 40MB for *S. aureus* to 437MB for combined human chromosome 14 dataset.

We find that Swalo can scaffold 19936 contigs from human chromosome 14 using 25.1 million reads in about 70 minutes and using 437MB of memory. This shows that it is efficient and scalable and thus applicable to scaffolding large genomes.

## Conclusions

The results show that Swalo performs consistently well and is able to identify many correct joins while keeping number of incorrect joins very low. It also shows pareto-optimal performance in the datasets we have analyzed *i.e.* there is a run of Swalo such that no other scaffolder in any of their runs was able to make more correct joins while making less than the number of incorrect links by Swalo. We observe that consistent results are achieved when Swalo is used with Bowtie 2. However, when reads are largely error free results achieved using Bowtie with no mismatches can be better possibly due to reads being mapped to more regions compared to Bowtie 2.

Overall we find that Swalo outperforms all other scaffolders on real and simulated datasets. This indicates that genome assembly may be substantially improved through the use of statistical models. The method may further be improved by modifying the heuristic used to select among multiple candidate joins and by considering global properties of the scaffold graph. The methods may also be extended to scaffolding with long reads generated by SMRT and nanopore sequencing. The improvement in scaffolding achieved by a practical method based on assembly likelihoods opens up the possibility that other problems related to genome assembly such as reference guided assembly, mis-assembly correction, copy number estimation, gap-filling may also be amenable to this approach.

## Materials and methods

We present here a brief description of methods behind Swalo. More detailed exposition is available in Supplementary Notes 3.1-3.4.

### Learning parameters and computing likelihood of contigs

The first step in Swalo is to estimate parameters and compute likelihood of contigs using the approach presented in [24] (Supplementary Note 3.1). The model incorporates insert size distribution, sequencing errors as well as randomness in read generation. We map reads to contigs and learn insert size distribution and error parameters using reads that map uniquely. The likelihood of contigs are computed with respect to a generative model using the learnt parameters. We use a smoothed and truncated version of the insert size distribution for scaffolding.

### Scaffold graph construction

We then construct the *scaffold graph* which is a bi-directed graph with a vertex for each contig and an edge between contigs if joining the contigs would lead to an increase in the assembly likelihood (Supplementary Note 3.2). The edges are weighted using this increase in likelihood. Edge weights are computed for each pair of contigs such that there are read pairs with two ends mapping to different contigs in the pair. This is done in two steps.

- First we estimate the maximum likelihood gap between the contigs using a generative model correcting for the issue that we may not observe inserts from the entire distribution. If there are read pairs that map to multiple pairs of contigs, we resolve them using the expectation maximization (EM) [27] algorithm.
- Then we check whether linking the contigs would result in an increase in the likelihood by computing probability of linking reads and adjusting probabilities of all other reads.

### Selecting joins

Once the scaffold graph is constructed, we first make unambiguous joins *i.e.* join contigs connected by an edge with an increase in likelihood and such that one vertex has outdegree one and the other has indegree one. The other possible joins are sorted according to the estimated increase in the likelihood and contigs are joined if likelihood increase of the candidate join is significantly higher than other conflicting joins as determined by a heuristic (Supplementary Note 3.3). If there are other joins consistent with the candidate join *i.e.* one or more contigs fit into the gap between the pair of contigs, we select from them using the dynamic programming algorithm for the weighted interval scheduling problem and remove conflicting ones. We choose a conservative approach during joining as unlinked contigs may later be merged using other datasets but incorrect joins would usually remain undetected for *de novo* assembly and might lead to errors in downstream analysis.

### Implementation

The methods have been implemented in a tool called ‘scaffolding with assembly likelihood optimization (Swalo)’ using C/C++. The read alignement and gap estimation phases are parallelized to speed up computation. Swalo is available for download freely at http://atifrahman.github.io/Swalo/.

### Data access

We use the data and analysis scripts used in [22] and [23]. Scripts to install tools, download data and generate the results used in this paper are available at https://github.com/atifrahman/Swalo/tree/scripts.

The scaffolds generated are also available at the same address. There may be minor variations in results due to thread race during mapping of reads unaligned by Bowtie as random subsets of unaligned reads are mapped. Original alignment files are available upon request.

## Acknowledgements

We thank Dan Rokhsar, Páll Melsted, Harold Pimentel, Shannon McCurdy and Nicolas Bray for helpful conversations during the development of Swalo. LP was funded in part by NIH R01 HG006129. AR was funded in part by Fulbright Science & Technology Fellowship 15093630.

## Author contributions

AR and LP conceived the project and developed the methodology. AR implemented the method in the Swalo software and obtained the results of the paper. AR and LP wrote the manuscript. All authors read and approved the final manuscript.

